# Evaluation of heating and chemical protocols for inactivating SARS-CoV-2

**DOI:** 10.1101/2020.04.11.036855

**Authors:** Boris Pastorino, Franck Touret, Magali Gilles, Xavier de Lamballerie, Remi N. Charrel

## Abstract

Clinical samples collected in COVID-19 patients are commonly manipulated in BSL-2 laboratories for diagnostic purpose. We used the French norm NF-EN-14476+A2 derived from the European standard EN-14885. To avoid the risk of exposure of laboratory workers, we showed that Triton-X100 must be added to guanidinium thiocyanate-lysis buffers to obtain a 6-log reduction of infectious virus. Although heating protocol consisting of 92°C-15min was more effective rather than 56°C-30min and 60°C-60min to achieve 6-log reduction, it is not amenable for molecular detection on respiratory specimens because of important decrease of detectable RNA copies in the treated sample *vs* untreated sample. The 56°C-30min and 60°C-60min should be used for inactivation of serum / plasma samples for serology because of the 5log10 reduction of infectivity and low viral loads in blood specimens.

## Introduction

Coronavirus disease 19 (COVID-19), classified as pandemic by WHO, is a severe acute respiratory syndrome (SARS) caused by the virus designated SARS-CoV-2 (1). Since December 2019, measures to reduce person- to-person transmission of COVID-19 have been implemented to attempt control of the outbreak. Tremendous efforts are done by an increasing number of scientific personnel working daily with the live virus and / or infectious samples, and thus heavily exposed to the risk of infection (2–4). Accordingly, the WHO introduced laboratory guidelines to mitigate this risk for diagnosis and research activities (5). Nonetheless, laboratory workers processing clinical samples will continue to be exposed to infectious SARS-CoV-2 (6). SARS-CoV-2 direct diagnosis is based on RNA detection by RT-qPCR (7). The methods for nucleic acid (NA) extraction use buffers, which formulation intends to obtain high quality NAs. They are not primarily developed for inactivation. Automated NA extraction is generally performed outside of biosafety cabinets which demands that only non-infectious samples must be loaded. To achieve this objective, a prior inactivation step under appropriate biosafety conditions is an absolute requirement. Previous studies have addressed the ability of lysis buffers added to the samples in initial step of NA extraction to act as inactivation agents of several pathogenic viruses (including coronaviruses). However, discrepant results observed with dissimilar protocols led to controversial conclusions (8–10). On another hand, the Center for Disease Control and Prevention (CDC) recommends using Triton X-100 and to heat the sample at 60°C for 1 hour for samples suspect of containing Viral Hemorrhagic Fever (VHF) agent. This procedure has been adopted by many laboratories for handling samples that may contain Ebola virus. Others studies with SARS-CoV and MERS-CoV have established that heat treatment can inactivate beta-coronaviruses (11,12). Consequently, definitive validation to SARS-CoV-2 is still awaited. Soon or later during the COVID-19 pandemic, serological tests will be used for diagnostics and for seroprevalence studies aiming at measuring the penetration of SARS-CoV-2 infection at population level. Detection of past infection will be pivotal for allowing immune persons to take back their professional activity. Since SARS-CoV-2 was detected in blood during infection (13), samples will have to be inactivated prior to serological tests are performed (14). In this study, we have tested ten different protocols including three lysis buffers and six heat inactivation procedures on SARS-CoV-2 culture supernatant.

## Materials and methods

### Lysis buffers

Three lysis buffers produced by Qiagen (Hilden, Germany) were tested. Approximate composition of each buffer is provided by Qiagen (18-20). ATL (1-10% sodium dodecyl sulfate [SDS]), VXL (30-50% guanidine hydrochloride, 1-10% t-Octylphenoxypolyethoxyethanol [Triton X-100]), and AVL (50-70% guanidinium thiocyanate). AVL has also been supplemented with 100% ethanol or 1% Triton X-100.

### Cell line

African green monkey kidney cells (Vero-E6; ATCC#CRL-1586) were grown at 37°C in 5% CO_2_ with 1% Penicillin/Streptomycin (PS; 5000U.mL-1 and 5000µg.mL-1; Life Technologies) and supplemented with 1% non-essential amino acids (Life Technologies) in Minimal Essential Medium (Life Technologies) with 5% FBS.

### Viruses

The Human 2019 SARS-CoV-2 strain (Ref-SKU: 026V-03883) was isolated at Charite University (Berlin, Germany) and obtained from the European Virus Archive catalog (EVA-GLOBAL H2020 project) (https://www.european-virus-archive.com). Experiments were performed in BSL3 facilities.

### SARS-CoV-2 titration

SARS-CoV-2 was first propagated and titrated on Vero-E6 cells. Virus stock was diluted to infect Vero-E6 cells at a MOI of 0.001; then cells were incubated at 37°C for 24-48 hours after which medium was changed and incubation was continued for 24 hr; then supernatant was collected, clarified by spinning at 1500 × g for 10 min, supplemented with 25mM HEPES (Sigma), and aliquoted. Aliquots were stored at - 80°C before titration. Virus infectivity was measured using 50% tissue culture infectivity dose (TCID_50_); briefly, when cells were at 80% confluence, six replicates were infected with 150μL of tenfold serial dilutions of the virus sample, and incubated for 3-5 days at 37°C under 5% CO_2_. CPE was read using an inverted microscope, and infectivity was expressed as TCID_50_/ml based on the Karber formula (15).

### Inactivation assays with lysis buffer (Table 1)

The French norm NF EN 14476+A2 derived from the European standard EN 14885 was used (16). For simulating “dirty” conditions, 3 g/L BSA was added before inactivation (Table 1). Each sample was incubated in duplicate with the lysis buffer at room temperature for 10 min; then lysis buffer was discarded via ultrafiltration with Vivaspin 500 columns (Sartorius, Göttingen, Germany) as described (17); column was washed with 500 µL PBS three times, and eluted in 20 µL of PBS; 0.1mL was inoculated onto Vero-E6 monolayer (70% confluence). Controls consisted of uninoculated Vero-E6 cells, Vero-E6 cells inoculated with the tested lysis buffer (cytotoxicity), and Vero-E6 cells inoculated with SARS-CoV-2 only. Cells were incubated at 37 °C under 5% CO_2_ for 5 days. The read-out was the presence of CPE together with SARS-CoV-2 RNA detection through RT-qPCR at day 5; in the absence of CPE at day 5, 100 µL of supernatant was passaged with the same read-out 5 days later (day 10).

**Table 1.** Protocols tested for assessing inactivation using lysis buffers.

| Lysis buffer | Composition<sup>a</sup> | Nucleic acid extraction kit (catalog #) | Interfering substance / added | Lysis buffer / sample | Temperature (°C) | Contact time (min) |
| --- | --- | --- | --- | --- | --- | --- |
| Buffer ATL | 1-10% SDS<sup>b</sup> | QIA Symphony DSP Virus/Pathogen Kits (#937036) or QIA Symphony DSP DNA Mini Kit (#937236) | ± BSA<sup>f</sup> (3g/L) | 1:1 | 20 | 10 |
| Buffer VXL | 30-50% GuHCl<sup>c</sup><br>2.5-10% Triton X-100<sup>d</sup> | QIA Amp cador Pathogen Mini kit (#54104) or QIA Amp 96 DNA QIAcube HT kit (#51331) | ± BSA (3g/L) | 1:1 | 20 | 10 |
| Buffer AVL | 50-70% GITC<sup>e</sup> | QIAamp Viral RNA Minikit (#52904) | ± BSA (3g/L) | 4:1 | 20 | 10 |
|  |  |  | ± BSA (3g/L) + 1 volume ethanol 100% | 4:1 | 20 | 10 |
|  |  |  | ± BSA (3g/L) + 1% Triton X-100<sup>g</sup> | 4:1 | 20 | 10 |

<sup>a</sup> as provided by Qiagen (18-20); <sup>d</sup> GuHCl, <sup>b</sup> Sodium dodecyl sulfate, <sup>c</sup> guanidine hydrochloride, <sup>d</sup> vol/vol, <sup>e</sup> guanidinium thiocyanate, <sup>f</sup> Bovine serum albumin, <sup>g</sup> final concentration (vol/vol).

### Heat inactivation assays (Table 2)

A 400-µL volume of SARS-CoV-2 supernatant (3.3×10^6^ TCID_50_ /mL) was incubated in a pre-warmed dry heat block and immediately tested for measuring TCID_50_ and RNA copies. Virus titration was performed in duplicate before and after heating to measure viral load reduction factor.

**Table 2.** Protocols tested for assessing heat inactivation.

| Sample tested | Interfering substance | Volume sample (μL) | Temperature ($^{\circ}$C) | Time (min) |
| --- | --- | --- | --- | --- |
| SARS-CoV-2 cell supernatant<br>($3.3 \pm 2.3 \times 10^6$ TCID<sub>50</sub>/ml)<sup>a</sup> | ± BSA<br>(3g/L)<sup>b</sup> | 400 | 56 | 30 |
|  |  |  | 60 | 60 |
|  |  |  | 92 | 15 |

### Integrity of SARS-CoV-2 RNA after heat inactivation

Heat inactivated samples were extracted using the Qiacube HT and the Cador pathogen extraction kit (both from Qiagen). Viral RNA was quantified by RT-qPCR (qRT-PCR EXPRESS One-Step Superscript™, ThermoFisher Scientific) (10min-50°C, 2 min-95°C, and 40 times 95°C-3 sec / 60°C-30 sec] using serial dilutions of a T7-generated synthetic RNA standard. Primers and probe target the N gene (Fw: GGCCGCAAATTGCACAAT; Rev : CCAATGCGCGACATTCC; Probe: FAM-CCCCCAGCGCTTCAGCGTTCT-BHQ1. The calculated limit of detection is 10 RNA copies per reaction.

## Results

### Inactivation assays with lysis buffer (Table 3)

VXL and ATL buffers were able to inactivate SARS-CoV-2 with viral loads as high as 10^6^ TCID_50_/ml. In contrast, AVL buffer (GITC 50-70%) either alone or in the presence of absolute ethanol or 1% Triton X-100 resulted in a partial inactivation (50-75%). In addition, our results show that GITC alone (AVL buffer) or GITC mixed with absolute ethanol also cannot guarantee SARS-CoV-2 inactivation as previously described (10). Finally, there was no difference observed between clean and dirty (3 g/L BSA) conditions.

**Table 3.** SARS CoV-2 inactivation using lysis buffers with additional reagents and with / without.

| Lysis buffer protocol | Virus detection (CPE + RT-qPCR) after inactivation |  |  |  |
| --- | --- | --- | --- | --- |
|  | Without BSA |  | With BSA(3g/L) |  |
|  | Replicate#1 | Replicate#2 | Replicate#1 | Replicate#2 |
| ATL buffer | No VR<sup>a</sup> | No VR | No VR | No VR |
| VXL buffer | No VR | No VR | No VR | No VR |
| AVL buffer | VR | VR | VR | VR |
| AVL buffer + 100% ethanol | VR | No VR | VR | No VR |
| AVL buffer + 1% Triton X-100 | VR | VR | VR | No VR |

<sup>a</sup> VR, SARS-CoV-2 replication; no VR defined as absence of CPE at passage#1 and passage#2 confirmed by RT-qPCR showing a Ct value >40; VR defined by CPE at passage#1 or passage#2 confirmed by RT-qPCR showing a Ct value <40.

### Heat inactivation assays (Table 4)

Only the 92°C-15min protocol was able to inactivate totally the virus (>6 Log_10_ decrease), whereas the two other protocols resulted in a clear drop of infectivity (5 Log_10_ reduction) but with remaining infectivity equal or lower than 10 TCID_50_/ml (Table 4). These results were consistent with previous studies on SARS-CoV and MERS-CoV (11,12). There was no difference between clean or dirty conditions.

**Table 4.**
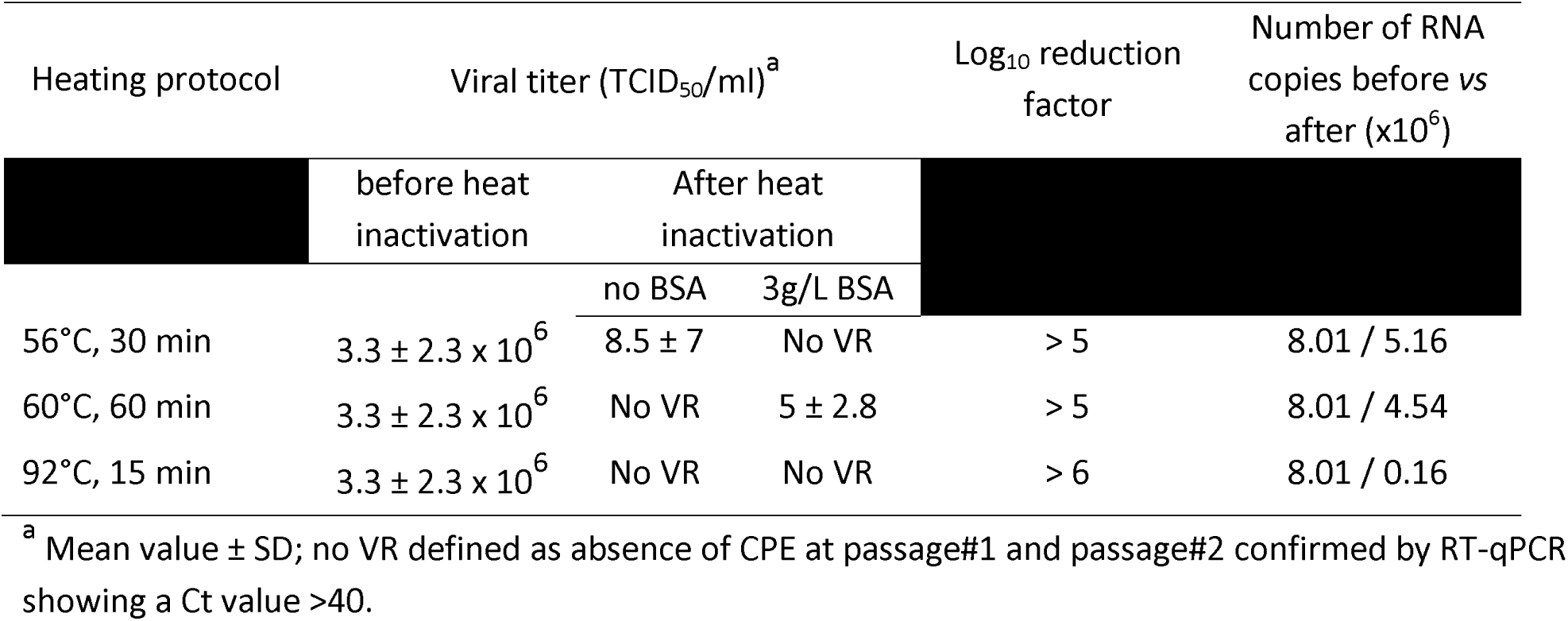
SARS-CoV-2 heat inactivation.

### Integrity of SARS-CoV-2 RNA after heat inactivation

The analysis of the Ct values (instead of the TCID_50_) showed that 56°C-30min and 60°C-60min did not affect significantly the number of detectable RNA copies (Δ Ct <1) (Table 4). In contrast, 92°C-15min resulted in a significant drop of the number of RNA copies (Δ Ct >5) (Table 4).

## Discussion

Despite the previous emergence of SARS and MERS CoV, there are few studies on the inactivation protocols aiming at mitigating the risk of exposure for medical and laboratory personnel (21).

Qiagen is a prominent actor in the field of nucleic acid purification. Most of other manufacturers of NA purification kits use similar lysis buffer as ATL, AVL and VXL. The ability of AVL to inactivate pathogenic viruses was debated (8–10) but there is no data for ATL and VXL. A total of ten different protocols using AVL, ATL and VXL alone or in association with ethanol or Triton-X100 were studied on SARS-CoV-2 according to the French version of the European recommended procedure (NF EN 14476+A2) (16), as previously shown for other viruses such as Ebola virus or Foot and Mouth Disease virus (8,10,22,23). Our results are in line with data reported for Zaire Ebolavirus (10). They strongly suggest that ATL or VXL should be preferred to AVL. Our findings corroborate and expand recent results (24).

Considering that low SARS-CoV-2 viremia is observed in COVID-19 patients even at the acute stage of the disease (21), the 56°C-30min and 60°C-60min protocols commonly used before serology appears as sufficient for inactivating SARS-CoV-2 as recommended before serological assay for other enveloped RNA viruses (25). Samples treated accordingly will also be amenable for viral RNA detection. In contrast, when processing respiratory samples commonly exhibiting much higher viral loads (26), only the 92°C-15min protocol showed total inactivation; however, whether this protocol is more efficient for inactivation than the two other, the drastic reduction of RNA copies that are detectable thereafter precludes its utilization for subsequent RT-qPCR detection of SARS-CoV-2. For the latter, inactivation using VXL, ATL or similar lysis buffer should be preferred.

Since clinical samples collected in COVID-19 suspect patients are commonly manipulated in BSL-2 laboratories, the results presented in this study should help to choose the best suited protocol for inactivation in order to prevent exposure of laboratory personnel in charge of direct and indirect detection of SARS-CoV-2 for diagnostic purpose.

